# Genome-wide expression gradient estimation based on local pseudotime in single cell RNA sequencing

**DOI:** 10.1101/2025.05.01.650773

**Authors:** Andreas Tjärnberg, Christopher A Jackson, Lionel A Christiaen, Richard Bonneau

**Affiliations:** Allen Institute for Brain Science, Seattle, Washington, USA; Center For Genomics and Systems Biology, New York University, New York, NY, USA; Sars International Centre for Marine Molecular Biology, University of Bergen, Bergen, Norway; Prescient Design a Genentech accelerator, New York, NY, USA

## Abstract

Single cell genomics measures the internal state of individual cells and makes it possible to describe biological phenomena such as cell type heterogeneity, developmental progression, and dynamic differentiation using computational methods. A prominent approach to facilitate downstream analysis, for large collection of cells, is to connect individual cells by a k-Nearest Neighbor Graph (kNN-G). The kNN-G is among other things used by methods that derive pseudotime, a temporal ordering of cells. However, pseudotime methods require knowledge of initial states of the trajectories created. Alternatives have been invented that estimate gene rate of transcription *e*.*g*. by RNA velocity but these methods don’t generalize to all individual genes for estimating transcriptional rate of change along pseudotime. While these methods can derive pseudo progression *de novo* by extrapolating transcription velocity vectors they are limited to subsets of genes where intronic reads are captured with sufficient magnitude. Here we present pseudovelocity, an alternative method for calculating a local rate of change in transcription based on the kNN-G and diffusion-based pseudotime, to derive RNA velocity estimates for individual genes, and demonstrate how it improves the gene-based estimation of RNA velocity for downstream analysis. This package can be accessed here: https://gitlab.com/Xparx/pseudovelocity.

## 1 Introduction

For single-cell transcriptomics data, individual cells can be connected through their similarity using the k-Nearest Neighbor Graph (kNN-G), a staple method used in the literature [1–3], which captures relationships in both the local and global cell transcription landscape. There are various efficient ways to calculate the kNN-G for a large number of cells, as well as using the kNN-G to project high dimensional data onto a manifold with dimension smaller than the original data [4, 5]. The resulting structure of the kNN-G can then be used to gain insight into the cellular landscape. One prominent example is to calculate the modularity and infer clusters of cells forming on the graph [6, 7] which can capture complex nonlinear relationships. The distance between cells estimated as a distance on the kNN-G can be interpreted under the right circumstances as a progression along a differentiation path. Pseudotime, a concept that is introduced as a cell ordering interpretation either on the graph or on a manifold projection of the cells, will be a natural interpretation if the cells have been collected throughout a dynamic process, such as during cell differentiation or over the cell cycle [[8–10]. In this setting, each cell would serve as a snapshot of a specific state representing a time point in its individual progression. The high abundance of measured cells can be connected through their transcriptional similarity, creating a complete time series [11–15].

A different method to estimate cell progression was described by La Manno et al. [8] which captures spliced and unspliced RNA from single cell experiments by creating a model of the ratio of spliced to unspliced counts and, given assumptions on the rate of splicing, showed that this could be used to estimate RNA-velocity (RNAv), *i*.*e*. a rate of transcription 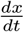 where *x* is expression and *t* is time. The assumption of rate of splicing is formulated as fitting a linear model of the ratio of normalized spliced to normalized unspliced mRNAs for each gene over all cells where the cells on the slope would give the steady state and deviation of the residuals would give rate of degradation *or* negative 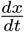, or active transcription, positive 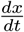. More specifically, steady state is given by the relationship *u* = *γs* where *u, s* are the unspliced and spliced transcripts respectivley and *γ* determines their steady-state relationship. In practice, the authors estimate *γ* from the lowest and highest percentile of data points. This choice is motivated by observed patterns of spliced–unspliced of the extreme ends of low and high expression. A revised approach was developed by [16] in which they infer reaction rates as part of estimating the dynamics along time as opposed to the steady state model used for estimating the original RNA velocity (RNAv) estimates. Conceptually, RNAv gives a natural way of estimating cell progression, by extrapolating a cell state from its estimate of the gene specific transcription rates. Then, by way of the cell similarity, one can project the direction of trascription towards neighbouring cells and determine a direction of cell development progression.

The “gold standard” would be to estimate RNAv from an individual cell with a reasonable normalization procedure; however, due to the nature of single cell expression RNAv suffers the same drawbacks as other single cell frameworks with the addition that there is a high risk of lowly sampled unspliced reads. The authors of RNAv report unspliced abundance upwards of 20 − 30% for their data where other observations in *e*.*g*. Wang et al. [9] have somewhere around ∼ 5% for all genes, dropping to ∼ 3% for the marker genes and genes of interest such as TFs (Figure 1B). With such low abundance any meaningful estimate would be overshadowed by noise and give a lower fraction of genes suitable for estimating RNAv, making RNAv of limited use for downstream analysis and gene-gene relationship inference. Coupled with single cell “noise” and low coverage and sample preparation variebility we would need to rely on imputing with the support of the cell neighborhood graph to average and boost the expression signal and the local neighborhood will be highly important to estimate the correct RNAv. RNAv is not decoupled from the neighborhood of cells and is therefore influenced by it. This is what in practice is done by La Manno et al. [8].

**Figure 1:**
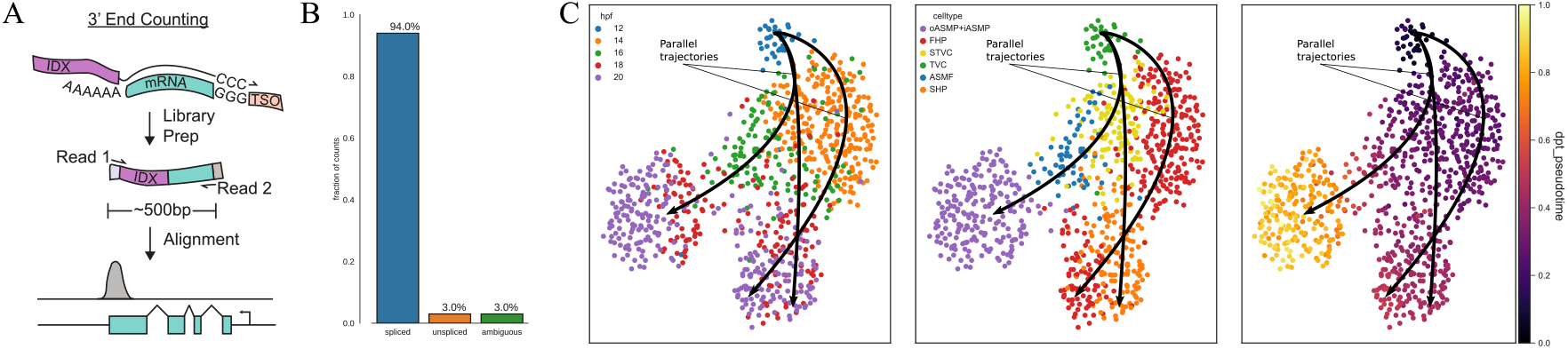
Aspects of single cell sequencing. A) Capturing mRNA using 10x and polyA barcoding. B) Spliced reads distribution in data collected by Wang et al. [9] using Drop-Seq2. C) TVC to heart cells similarity when acquiring specified fates[9].

Some experimental techniques are less likely to capture unspliced read, such as the popular 10X experimental framework[17] where there is a transcript end bias for captured reads, Figure 1A, significantly lowering the coverage of RNAv to a small subset of genes where the intronic reads are able to be captured. For the purpose of trajectory inference this might be sufficient. But that is not the case for GRNi.

## 2 Results

Using gene expression in general we would expect that the RNAv estimates would correlate with the rate of change in the total amount of mRNA along the expression trajectory. This is explored briefly by the authors of RNAv[8]. Here we compare the rate of change in mRNA on the local neighborhood to estimated gene rate of change, pseudovelocity, (PV) and compare to the estimate of RNAv derived by scVelo[16]. The advantage of using RNAv would be that we could in theory estimate RNAv for each cell independently, but, as we have seen, the estimate of RNAv currently relies heavily on the estimate of *γ* and kNN imputation on the cell neighborhood. Here we take advantage of the cell-cell relation to estimate an RNA “velocity” based on distribution of mRNA over the neighborhood.

A motivation for using principle curves and rates in downstream analysis is to estimate cell ordering and projecting all cells onto this pseudotime trajectory. Projecting the cell order onto one trajectory has been shown to perform poorly compared to using actual transcription rates for the basis of GRN inference[18]. As outlined in section 4.1, 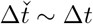 only if Δ is sufficiently small. If the cells perpendicular to the trajectory are projected on to the principle curve this assumption may not hold and we would estimate highly noisy values of 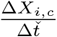 with possibly different regulatory programs active for neighboring cells. This relationship can be seen for the first and second heart progenitor fate trajectories in *C. robusta* for cells that have divided (FHP and STVC) but are aligned closely in the umap space (figure: 1 panel B)[9]. It is also the case that the principle curve is dependent on the projection used to calculate it, making it break the proportionality constraints detailed above. Papili Gao et al. [19] uses a discretized model to infer their system, which disregards the continuous nature of cell progression and requires time stamped data or assumptions about delta times to work.

It should also be noted that the definition of RNAv used in works such as Qiu et al. [18] assumes that the RNAv presented by La Manno et al. [8] is conceptually the approximation of 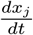 and can be used to explain the magnitude of *x*, which is not the case (Figure 2C, D, and 3D).

**Figure 2:**
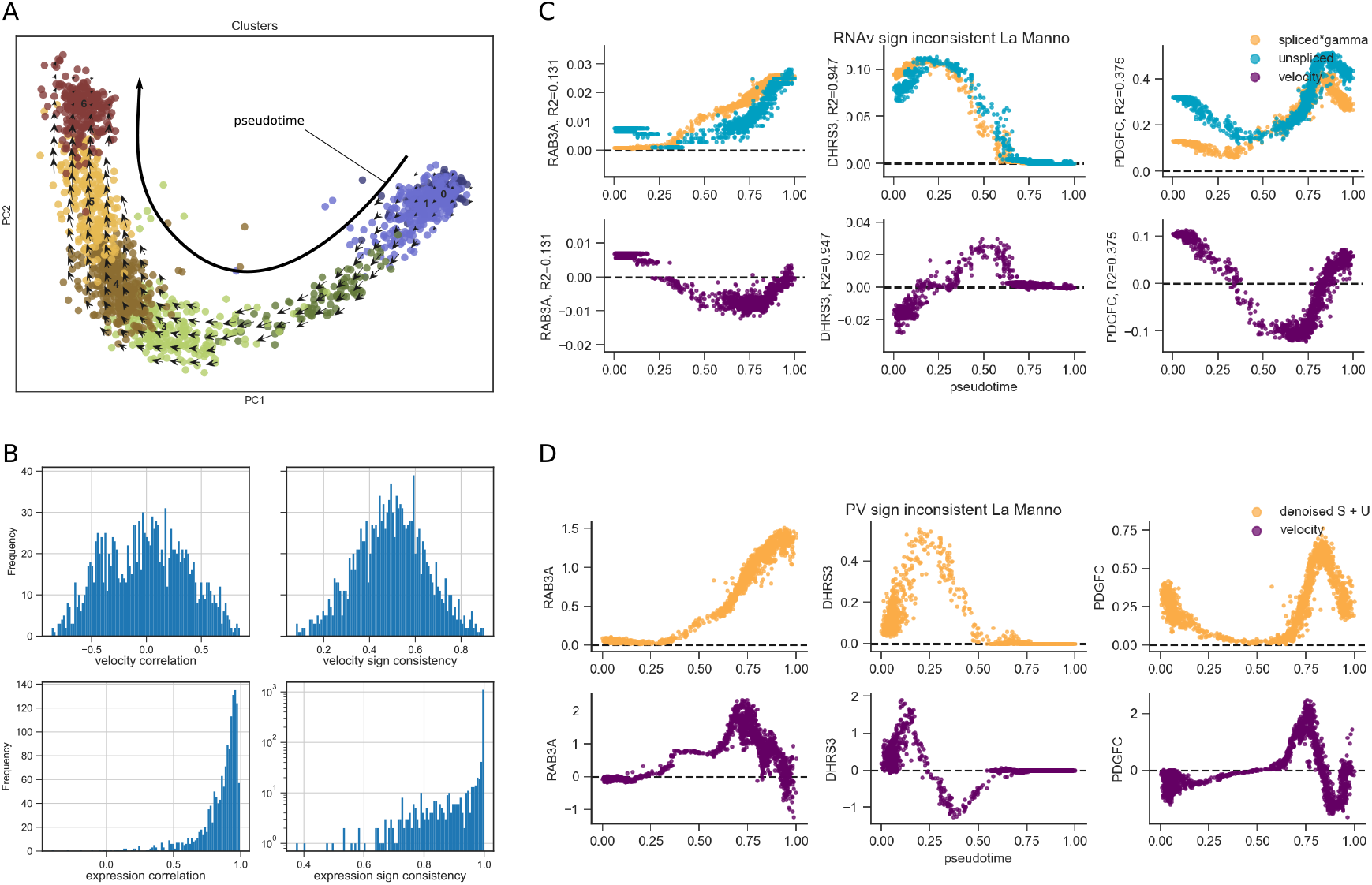
*Pseudovelocity* (PV) compared to RNA-velocity (RNAv) for the hgForebrainGlut [8]. A) Reproduced trajectory and PV projection in two first principle components. B) Genome wide agreement, after filtering out genes removed by RNA-velocity preprocessing, of expression profiles and velocity profiles between RNAv and PV. Velocity linear correlation between RNAv and PV for all genes (upper left) with corresponding expression correlation (lower left) between preprocessed expression values. Sign overlap {−1, 0, 1} between RNAv and PV (upper right) and corresponding expression sign (lower right). Discrepancy in expression sign is due to zeroed value genes for some cells between the different normalization. C) 3 most sign -inconsistent genes between RNAv and PV expression profiles (spliced *× γ* and unspliced, top row), and corresponding RNAv (bottom row). D) 3 most sign-inconsistent gene expression profiles for preprocessed expression (top row) and PV (bottom row).

**Figure 3:**
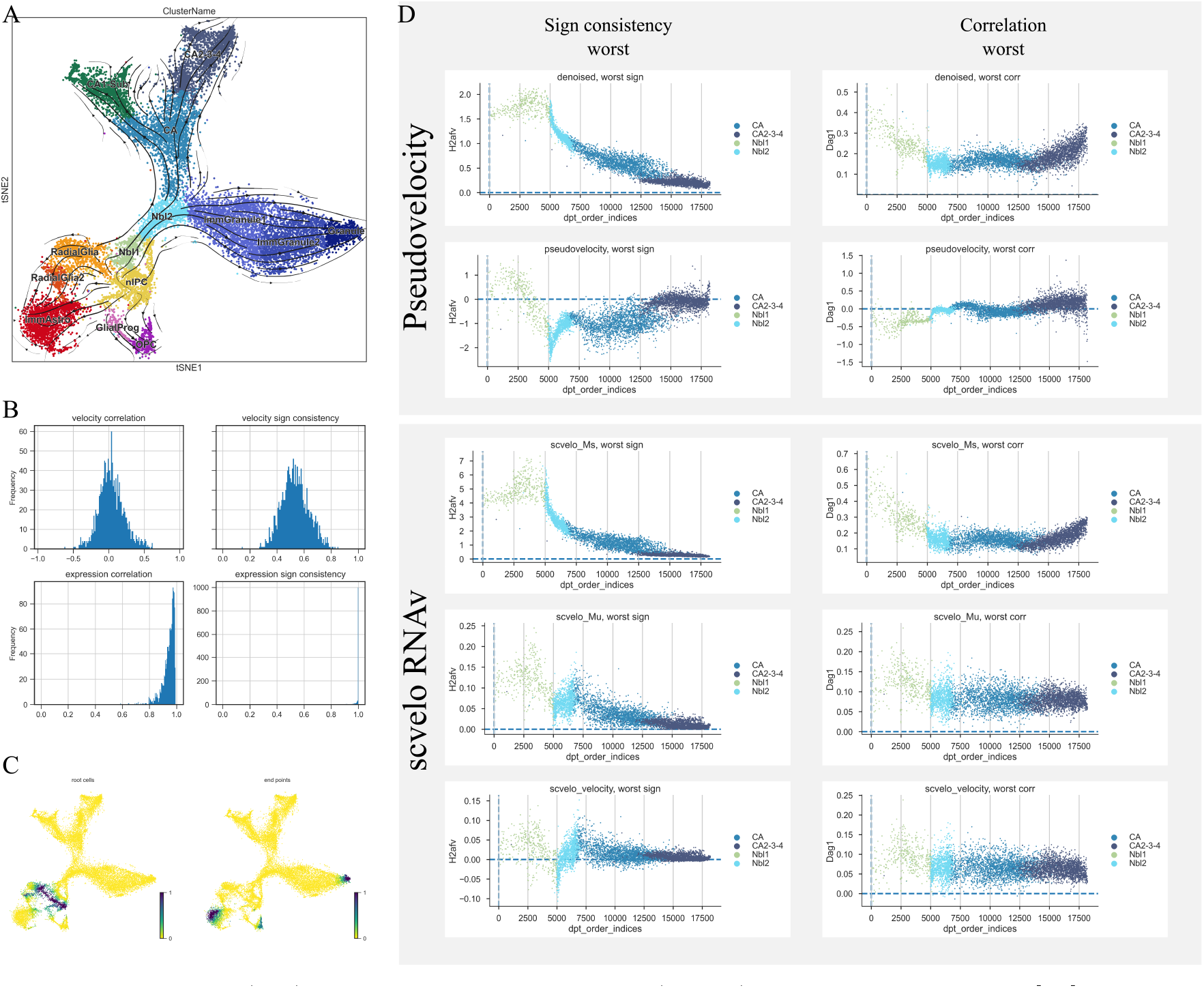
*Pseudovelocity* (PV) compared to RNA-velocity (RNAv) computed with scvelo [16] for the dentate gyrus [8]. A) tSNE projection computed by La Manno et al. [8] with PV projected velocity estimates projected on top. B) Genome wide agreement, after filtering out genes removed by RNA-velocity preprocessing of Expression profiles and velocity profiles between scVelo RNAv and PV. Velocity linear correlation between RNAv and PV for all genes (upper left) with corresponding expression correlation (lower left) between preprocessed expression values. Sign overlap {−1, 0, 1} between RNAv and PV (upper right) and corresponding expression sign (lower right). Discrepancy in expression sign is due to zeroed value genes for some cells between the different preprocessing steps. C) Inferred root cells and end of trajectory cells marked by scVelo using PV. D) The genes H2afv and Dag1 showing the worst sign consistent and correlated genes between PV and scVelo velocity estimate. For PV the expression profile (top) and velocity estimate (bottom) is shown. For scVelo the mean spliced (Ms/top), unspliced (Mu/middle) and velocity (bottom) is shown.

We also explore an approach to infer a maximal decay rate for different cell contexts using the methods outlined in section 4.1.1. For this experiment we use the cell cycle in *S. cerevisiae* grown in an optimal nutrient environment (YPD)[10]. By labeling the cell cycle phases by marker genes we can trace a dynamic transcriptional process mapping the cell cycle, Figure 4A. The decay rate is a gene centric feature that can be driven by active decay factors but can not be part of the process of transcription and is therefore a crucial component to incorporate in GRN models that explain transcription by transcription factors (TFs). We look at two cell cycle dependent genes PIR1 and MRH1. These genes are active in the cell cycle in different phases, PIR1 activated in M=phase and peaking in M-G1 and deactivated in late M-G1 to G1. MRH1 is turned on in G2-phase peaking in M-phase, Figure 4B. Decay rates can then be computed by computing the dependency between observed expression levels and inferred rates of changes as described in section 4.1.1. This experiment is shown in figure 4C. We demonstrate that the decay rate can be set to a maximal bound, here divided into each cell cycle phase, as a liner factor dependent on the amount of transcripts as a function of the estimated rate of change.

**Figure 4:**
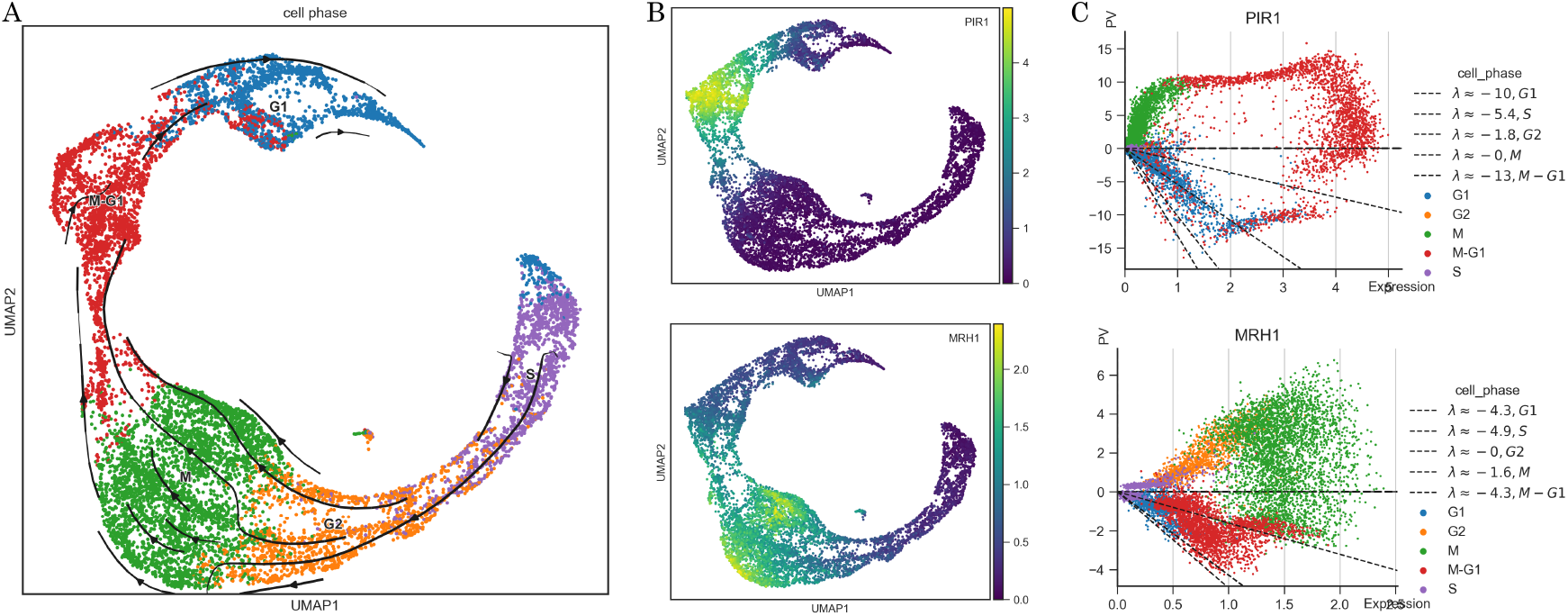
*Pseudovelocity* for circular dynamic of the *Saccharomyces cerevisiae* cell cycle and phase specific decay rate estimates[10]. A) UMAP projection of the cell cycle in *S. cerevisiae* in rich growth medium. Pseudo velocity direction of dynamics projected on top. B) UMAPs showing the denoised[20] expression level of PIR1 and MRH1. C) Inferred PVs plotted against expression level with decay rates indicated as the lowest negative slope capturing the negative PVs within the 5th percentile lowest value. A different decay rate was computed for each cell phase in the cell cycle.

## 3 Discussion

In this work we demonstrate that it is feasible to derive a gene wise rate of RNA production (RNAv) by regressing the local cell neighborhood transcription profile against diffusion pseudotime (pseduovelocity). This is demonstrated using single cell benchmark datasets derived for this specific purpose and are shown in ideal conditions. We show that the gene wise estimates derived from using intron and exon reads does not always estimate an RNAv that coincide with the rate of change in total mRNA abundance. While the biological underpinning for this might have a biological foundation, *e*.*g*. the rate of splicing and degradation rates might differ from the assumptions made in these methods, it is shown that even core properties like the sign determining the direction of change can be significantly different from what is observed. For the purpose of deriving GRNs and other downstream tasks, these assumptions need to be accounted for. Pseudovelocity require a predefined direction of cell development derived from methods like diffusion pseudotime while RNAv derived from intron and exon reads *de novo* infer direction of transcription. A future method that use intron–exon RNAv as a basis for pseudovelocity can be derived for re-estimating RNAv estimates across the trajectory for all genes in the transcriptome. We also demonstrate that, by setting up an ODE model of transcrption with separate mRNA decay and production rates, we can estimate a bound for the decay component further aiding in disentangling effects in the transcriptional machinery and make it possible to create a biophysical model for inference of regulation.

We propose pseudovelocity as an alternative estimate of RNAv that can be used for downstream analysis of gene wise transcription rate of change for analysis related to transcriptional behavior across development and other dynamic cellular processes.

## Methods

### Local gradient regression (pseudovelocity)

We want to estimate the rate of transcription

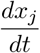

where *x*_*j*_ is the mRNA expression of gene *j* at time *t* in some unit. Each cell can be viewed as a snapshot in time relative to the internal process of the cell and can be globally related to the total process by the cell-cell relationships through the concept of pseudotime 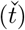. The cell specific time will be synced based on cell similarities and will not represent wall clock time but rather the cell internal process time, see [12] supplementary notes for a thorough walk-through of the concept of universal time and pseudotime.

For very densely collected data we would expect that 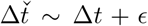 for pairs of cells if Δ is sufficiently small and represents a measurement where cells change in time. Given these assumptions, the difference between delta 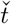 and delta *t* would vanish for sufficiently small deltas. With a somewhat relaxed assumption of linearity close to Δ ∼ 0, 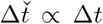. Therefore, an estimate 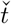 that is standardized over the complete cell population with a constant proportionality factor that does not vary depending on the neighborhood is preferred. Haghverdi et al. [12] derives the pseudotime measure *diffusion pseudotime* (dpt) which normalizes the Markov diffusion pseudotime metric over all path lengths on the diffusion map and would be one of the more suitable estimates of pseudotime to use to estimate

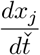

for this purpose. This gives a 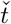 that is unitless and 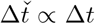 as 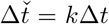 where k maps universal time to our scaled pseudotime.

We can reformulate the above rate to

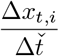

given the argument above about proportionality we can apply the chain rules so that

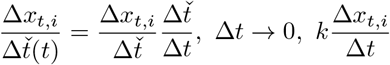

with *k* being the relation between time scales. If we can linearize expression changes close to the cell of interest we can estimate the rate of change in mRNA expression over closely localized cells with

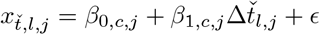

where *j* is the gene and *l* is the cells in the neighborhood of cell *c, l* ∈ *N*_*c*_. 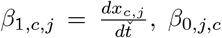 is the expression of gene *j* at the cell relative time 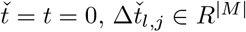 is the difference in pseudotime from cell *l* to cell *c* and 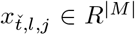 is the expression of the gene *j* in cell *l* at time 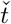. We could in theory subtract 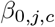 from 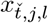 and not estimate it, however this would make us more susceptible to noise and we would not be able to estimate a rate when gene *j* is a drop-out at 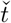 and has 0 expression, which we might want to do.

#### 4.1.1 Inferring decay rates

If we use an ODE model to model transcription of a gene *j* we know that transcription from TFs regulate the speed of mRNA production while the decay of those mRNAs are controlled by the stability of the mRNA itself as well as the decay factors able to degrade mRNA so that the rate of mRNA production in the cell;

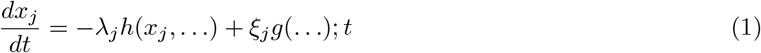

Here we introduce two functions *h*(…) and *g*(…) separately regulating two components, the decay of mRNA in a cell, where *h* is also dependent on the amount of transcripts of gene *j*. If we limit the mRNA regulatory effects to the decay rate of the gene itself, we seek the maximum degradation rate for each gene, given an estimate of 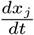 and the constraints *λ*_*j*_ *>* 0 and *ξ*_*j*_ ≥ 0 (equation 1) we can pose the following problem,

In matrix notation, let **Λ** ∈ *R*^*n,n*^ be the diagonal matrix with **Λ**_*i*=*j*_ = *λ*_*j*_ and **Λ**_*i* ≠*j*_ = 0 then

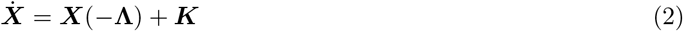

where 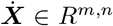 is the rate of all genes *n* in all samples *m*, ***X*** ∈ *R*^*m,n*^ ≥ 0 is the expression matrix. ***K*** is the rate of 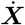 not explained by the decay rate which contains the transcription rates *g*(.).

Our aim is to solve for **Λ**;

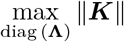

subject to:

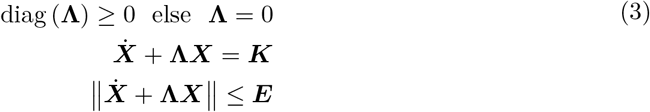

By noting that each gene can be solved independently due to the the fact that **Λ** is diagonal we can also write the gene-wise solution as

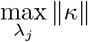

subject to:

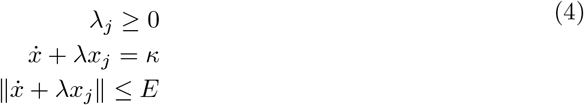

The objective max_**Λ**_ ||***K***|| would give the estimates with the maximum negative gene-wise bound of *λ*_*j*_. However, in our problem setup we have to introduce ***E*** to bound the solution to a reasonable allowed value. How to choose this penalty optimally has to be found.

We need to choose ***E*** to be able to solve for the objective. In practice we need to find a more heuristic way of solving the optimization 4 if we can’t set ***E***.

We choose to compute *λ*_*j,l*_ for each gene *j* and cell *l*, by looking at the maximum negative fraction

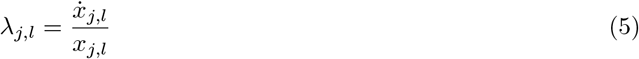

Choosing *λ*_*j,l*_ as the smallest rate among the lowest 5’th percentile of rates 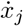. This means that we pick the slope of decay as the largest slope for the 95’th percentile cells.

